# Xolographic printing of cell-dense engineered living matter via iodixanol-driven enhancement of type II photoinitiation

**DOI:** 10.64898/2026.09.29.755253

**Authors:** Annalise Anspach, Alexis Wolfel, Mariel Cano-Jorge, Floris Dalenoord, Niklas Felix König, Catarina A. Custódio, Stan Rolfes, João F. Mano, Robert Passier, Jeroen Leijten

**Affiliations:** Leijten Lab, BioEngineering Technologies, TechMed Centre, Faculty of Science and Technology, University of Twente, 7522NB, Enschede, The Netherlands; Department of Applied Stem Cell Technologies, BioEngineering Technologies, Cardiovascular Health Technology Centre, TechMed Centre, Faculty of Science and Technology, University of Twente, 7522NB, Enschede, The Netherlands; xolo GmbH, Volmerstraße 9B, 12489 Berlin, Germany; CICECO – Aveiro Institute of Materials, Department of Chemistry, University of Aveiro, 3810-193 Aveiro, Portugal; Metatissue, PCI - Creative Science Park, Via do Conhecimento, 3830-352 Ílhavo, Portugal

**Keywords:** Refractive index, Bioxolography, volumetric bioprinting, photorheology, type II photoinitiators, cell-dense

## Abstract

Type II photoinitiators are essential for numerous applications, including advanced 3D printing techniques such as Xolography. Despite offering tunable wavelengths and reaction mechanisms, their limited photoreactivity hinders widespread adoption. Solutions to increase reactivity, particularly in aqueous solutions and in a biocompatible manner, are urgently needed. Here, we report the surprising finding that iodixanol enhances reactivity of type II photoinitiation systems. Supplementing Xolographic print resins with iodixanol increases photoreactivity, which reduces print times and light dosage tenfold, and enables printing of previously unprintable material formulations. Surprisingly, iodixanol supplementation also improves cell viability in a dose-dependent manner, while reducing cell-mediated light scattering. Together, this enables accurate Xolographic printing of highly cell-dense bioresins, which facilitated the biofabrication of engineered living matter with up to 100 M cells mL^-1^ (>40% cells by volume) with a resolution of ∼65 µm. These advances enabled the first demonstration of tissue function in Xolographically-printed materials. Human induced pluripotent stem cell-derived cardiomyocytes were printed into contractile cardiac-like engineered living matter, which exhibited enhanced cell-cell contact and synchronous contractions, confirming tissue level function. As a highly water soluble, cytocompatible, and clinically approved compound, iodixanol therefore represents a valuable and translatable solution to improve the reactivity of type II photoinitiation systems, which enables volumetric biofabrication of highly cell dense living matter. This unlocks a new role for iodixanol in a wide variety of fields including dentistry, chemistry, organic photovoltaics, tissue engineering, and 3D printing.

## Introduction

Photopolymerization is of indispensable value for a wide variety of fields, including dentistry^[1]^, chemistry^[2]^, organic photovoltaics^[3]^, tissue engineering^[4]^, and 3D printing^[5]^. Type I photoinitiators have recently been the focus of research and product development owing to their ability to crosslink with straightforward chemistry and fast reaction rates. However, much historical and current research has focused on leveraging the potential of type II photoinitiators, as they offer greater ranges of absorption wavelengths and reaction mechanisms.^[6]^ For example, peptides and other functional materials may act as coinitiators in type II photoinitiation systems, which offers the benefit of encoding the construct with additional functionality while preventing leaching, as the coinitiator becomes part of the final matrix.^[7]^ Moreover, the chemical versatility and broad range of absorption wavelengths of Type II PIs is instrumental to developing novel technologies. Recently, Xolography, a next-generation volumetric 3D printing technology, has been achieved using type II photoinitiators that mechanistically respond to irradiation with two different but simultaneous light wavelengths.^[8]^ However, the low photoreaction rate of type II photoinitiation systems limits its application as it demands higher light doses and longer exposure times to achieve crosslinking. Novel solutions to improve the reaction rates of type II photoinitiation systems are therefore urgently needed.

Various strategies to increase type II photoreactivity have been explored, most commonly via the addition of iodonium salts.^[9, 10]^ However, due to their low water solubility and high cytotoxicity, these are of limited use for aqueous and biomedical applications. Furthermore, although many type II photoinitiators themselves are highly cytocompatible, available coinitiators are typically cytotoxic.^[11–13]^ As Xolography relies on a dual-color type II photoinitiator, these issues are critical, particularly for the fabrication of engineered living matter.

In Xolography, a dual-color photoinitiator (DCPI) enables the light-sheet based printing technique to crosslink photopolymerizable materials specifically where two light beams of different wavelengths intersect. This approach enables the printing of centimeter-scale objects with 10 µm resolution within mere minutes.^[8]^ Furthermore, this technique allows for precise spatial encoding of mechanical properties and molecular patterns.^[14]^ Although the iodonium salt diphenyliodonium chloride (DPI) was recently shown to facilitate cytocompatible printing of low cell density cartilage-like living matter, the use of Xolographic printing for engineering tissues has remained limited. Specifically, the low photoreaction rate has restricted Xolographic (bio)printing to high concentrations of highly reactive materials, while only allowing for the use of low concentrations of highly resilient cell types, such as human mesenchymal stem cells and human primary chondrocytes exhibiting only cell level, rather than tissue level function.^[12–14]^ Indeed, all reported volumetric bioprinting techniques have proven unable to print cell-dense bioresins, with Xolography reaching maximally 1 M cells mL^-1[13–15]^ and computed axial lithography 15 M cells mL^-1[16]^, equating to cells constituting less than 0.5% and 5% of print volume, respectively. Not surprisingly, the inability to volumetrically print at high cell density has historically challenged a wide variety of its biofabrication applications as it hinders realization of tissue level organization and associated function. We reasoned that the linear nature of Xolography could result in reduced sensitivity to light scattering compared to computed axial lithography, thereby making Xolography particularly suitable for high cell density printing. However, current type II photoreaction rates remain too low for adequate crosslinking in the presence of high cell densities. Additionally, low reactivity severely limits the material toolbox that Xolography could access for the (bio)fabrication of soft matter, hindering its versatility and its widespread use in various applications such as soft robotics, wearable sensors, and tissue engineering.

Here, we report the surprising finding that iodixanol can be used as a single additive to drastically improve the reactivity of type II photoinitiation systems. This photoreactivity increase greatly reduced required energy dose and total print time for Xolographic prints. Additionally, this allowed for reduction in the required amount of coinitiator, which improved cytocompatibility and, surprisingly, further increased photoreactivity. Moreover, we show that the increased reactivity enables printing with a wider range of materials, including those with lower reactivities, such as human protein-based bioresins. As an added benefit, iodixanol can reduce scattering via refractive index matching^[16, 17]^, which in combination with higher reactivity enabled for the first time Xolographic printing of highly cell-dense (100 M cells mL^-1^, >40% cells by volume) living matter. This enabled rapid, high-resolution Xolographic printing of highly sensitive human induced pluripotent stem cell-derived cardiomyocytes (hiPSC-CMs) into biofabricated contractile living matter. This simple yet novel approach has significant implications for the rapid, high-resolution, and cytocompatible biofabrication of highly cell-dense living matter using volumetric printing techniques. Furthermore, although the ability of iodixanol to boost type II photoinitiation is unreported, iodixanol is a well-characterized, clinically approved^[18]^, highly water soluble, iso-osmotic, non-ionic, and cytocompatible compound^[19]^. This positions iodixanol as an enabling solution for type II photoinitiation in a wide variety of fields including dentistry, chemistry, organic photovoltaics, tissue engineering, and 3D printing.

## Results and Discussion

### Iodixanol improves reactivity of type II photoinitiation systems

Bioxolography is a light sheet-based volumetric bioprinting technique that uses DCPI to crosslink cell-laden photopolymerizable materials where two distinct yet specific wavelengths of light intersect. (**Figure 1a**). However, the low photoreactivity of current DCPIs in aqueous media challenges the Xolographic printing of hydrogels. It was recently demonstrated that DPI improves the reactivity of type II DCPI systems, where DPI oxidizes activated DCPI back into its original state, increasing the rate at which free radicals are formed.^[14]^ However, despite its ability to strongly improve reactivity, DPI’s low water solubility and cytotoxicity limits its use in tissue engineering. We investigated IDX with the aim of increasing the refractive index of the bioresins, but we consistently observed reactivity enhancements that suggested a broader functional role. Motivated by structural analogies between DPI and IDX, particularly the presence of iodine-substituted aromatic rings, we hypothesized that iodixanol could serve as a biocompatible additive and alternative to DPI and other iodonium salts (**Figure 1b**). We confirmed iodixanol’s high water solubility (75 wt%) at >20 fold more than DPI’s (**Figure 1c**). Furthermore, we demonstrated that while DPI exhibited cytotoxicity on a 3T3 fibroblast monolayer at concentrations as low as 1.6 wt%, iodixanol showed no obvious cytotoxicity, even at concentrations as high as 34 wt% (**Figure 1d**). Although this confirmed that iodixanol is highly compatible with the formulation of bioresins, the potential effect of iodixanol on type II photopolymerization has never before been studied (**Figure 1e**).

**Figure 1.**
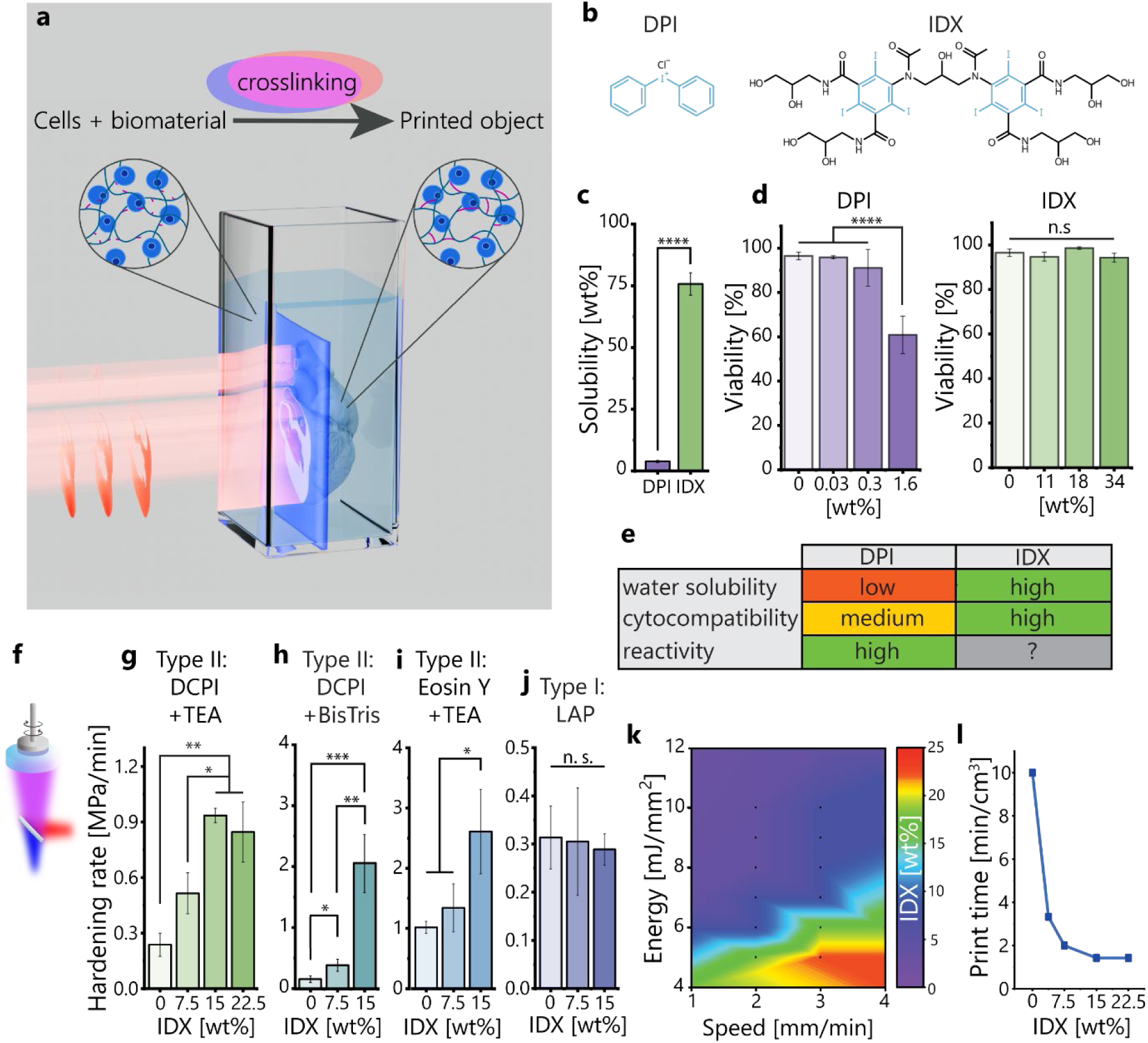
Iodixanol improves reactivity of DCPI-based hydrogel print formulations. (a) Schematic depiction of Bioxolography, where a UV light sheet activates the DCPI to the latent state, while a visible light video of a sliced object is projected on the cuvette to activate the DCPI. Where the light sheet and visible light intersect, crosslinking occurs, building up an object. (b) Chemical structures of DPI and iodixanol (IDX). (c) Solubility of DPI and iodixanol in water (n=3). Analysis is a two sample t-test. (d) Viability of 3T3 fibroblast monolayer incubated with DPI and iodixanol (n=3). (e) Summary of relevant properties of DPI and iodixanol. (f) Schematic representation of dual-color photorheology setup. (g) Effect of iodixanol on the hardening rate of 5% GelMA containing DCPI+TEA, (h) 5% GelMA with DCPI+BisTris, (i) 10% PEGDA with eosin Y+TEA, or (j) LAP as photoinitiation systems (n=3). (k) Iodixanol concentration that gives best quality prints (Figure S1) for different print speed-energy combinations. (l) Effect of iodixanol concentration on the time it takes to print a 1 cm^3^ object. Data is shown as mean ± standard deviation. Analysis is a one-way ANOVA with a post-hoc Tukey test. ****p<0.0001

To explore whether iodixanol boosts photoreactivity, we developed a dual-color photorheological setup to simulate Xolographic printing and investigate photocrosslinking (**Figure 1f**). The speed at which the formulation stiffness increased (i.e., hardening rate) is proportional to the reactivity (i.e., a more reactive formulation should crosslink and therefore harden more rapidly). This accounts for the premeasurement thermogelation of the gelatin methacryloyl (GelMA) based print formulations at printing conditions (i.e., room temperature). Dual-color photorheology revealed that iodixanol increases the hardening speed in a dose-dependent manner up to ∼4 fold compared to pristine formulations using TEA as a coinitiator (**Figure 1g**), or >10 fold when BisTris is used as a coinitiator (**Figure 1h**). To investigate whether this increase in photoreactivity was specific for DCPIs or applicable to other photoinitiators, the effect of iodixanol on hardening speed was investigated using eosin Y, methylene blue, and riboflavin, all type II photoinitiators used in light-based 3D printing. For each of these distinct type II photoinitiator systems, a dose-dependent increase in crosslinking rate was demonstrated (**Figure 1i**, **Figure S2m-n**). Formulations containing eosin Y also demonstrated a dose-dependent photocrosslinkability increase (**Figure 1i**). In contrast, iodixanol had no effect on type I photoinitiators such as lithium phenyl-2,4,6-trimethylbenzoylphosphinate (LAP), suggesting that iodixanol specifically improves type II photoinitiation systems (**Figure 1j**). Moreover, UV-vis experiments on the effect of iodixanol on poly(ethylene glycol) diacrylate (PEGDA) polymerization confirmed that iodixanol increases reactivity, yet also revealed that no crosslinking occurs in the DCPI-iodixanol system without the presence of a coinitiator (**Figure S3**). This confirms that iodixanol does not act as a coinitiator and suggests that iodixanol may act synergistically with the DCPI and coinitiator to increase reactivity of the DCPI-iodixanol-coinitiator system. Furthermore, while iodixanol-driven reactivity increase appears broadly applicable to type II photoinitiators, it may be specific to reductive coinitiators (i.e., TEA/BisTris), as no increase in photoreactive rate was observed for oxidative coinitiators (i.e., sodium persulfate) (**Figure S2o**). Of note, DPI exhibits similar behavior, showing no improvement in hardening rate when a oxidative coinitiator is used (**Figure S2p**). Together, this identifies iodixanol as an additive and alternative to iodonium salts to boost type II photoreactivity.

To determine iodixanol’s effect on printing conditions, we systematically varied the iodixanol concentration in the print resin and Xolographically printed at different speeds and UV light energies. This yielded a comprehensive map of the ideal iodixanol concentration for each print condition (**Figure 1k**). Generally, iodixanol enabled printing with both lower energy and higher speed in a dose-dependent manner. This is particularly desirable for cell-containing printing, as it decreases the UV dose cells receive from the light sheet, reducing potential phototoxicity and time spent outside of ideal culture conditions. Of note, even small amounts (3.75 wt%) of iodixanol allowed for a large (70%) reduction in print energy. Similarly, iodixanol exerted a dose-dependent effect on print speed, meaning that total print time could be reduced up to 85% via iodixanol addition (**Figure 1l**). This uniquely allows Xolographic printing of hydrogels to achieve similar production speeds to other volumetric printing techniques, such as computed axial lithography.^[20]^ As Xolography is positioned to scale up production of large living models, it is anticipated that reducing print times will be critical to its success and widespread adoption.

### Iodixanol enables improved cytocompatibility of bioresins

Type II photoinitiation demands the use of a coinitiator, such as triethanolamine (TEA), which acts as an electron donor to trigger photopolymerization. Unfortunately, practical concentrations of TEA for efficient bioprinting, including Bioxolography, causes cell death.^[14, 21, 22]^ We hypothesized that an iodixanol-driven increase in photoreactivity would allow for reduction of TEA concentration while maintaining effective photopolymerization. To this end, we investigated the effect of varying TEA and iodixanol concentration on the photopolymerization of 5% GelMA using dual-color photorheology (**Figure 2a**). In the absence of iodixanol, TEA concentration positively correlated with hardening rate, as expected. Surprisingly, when iodixanol was added, lowering the TEA concentration yielded drastically higher photoreactivity. Notably, the highest level of photoreactivity was achieved with bioresins containing 1 wt% TEA and an iodixanol concentration between 15 wt% and 22.5 wt%. Here, the addition of iodixanol allowed for a fivefold reduction in TEA concentration, while simultaneously achieving a doubling of the hardening rate.

**Figure 2.**
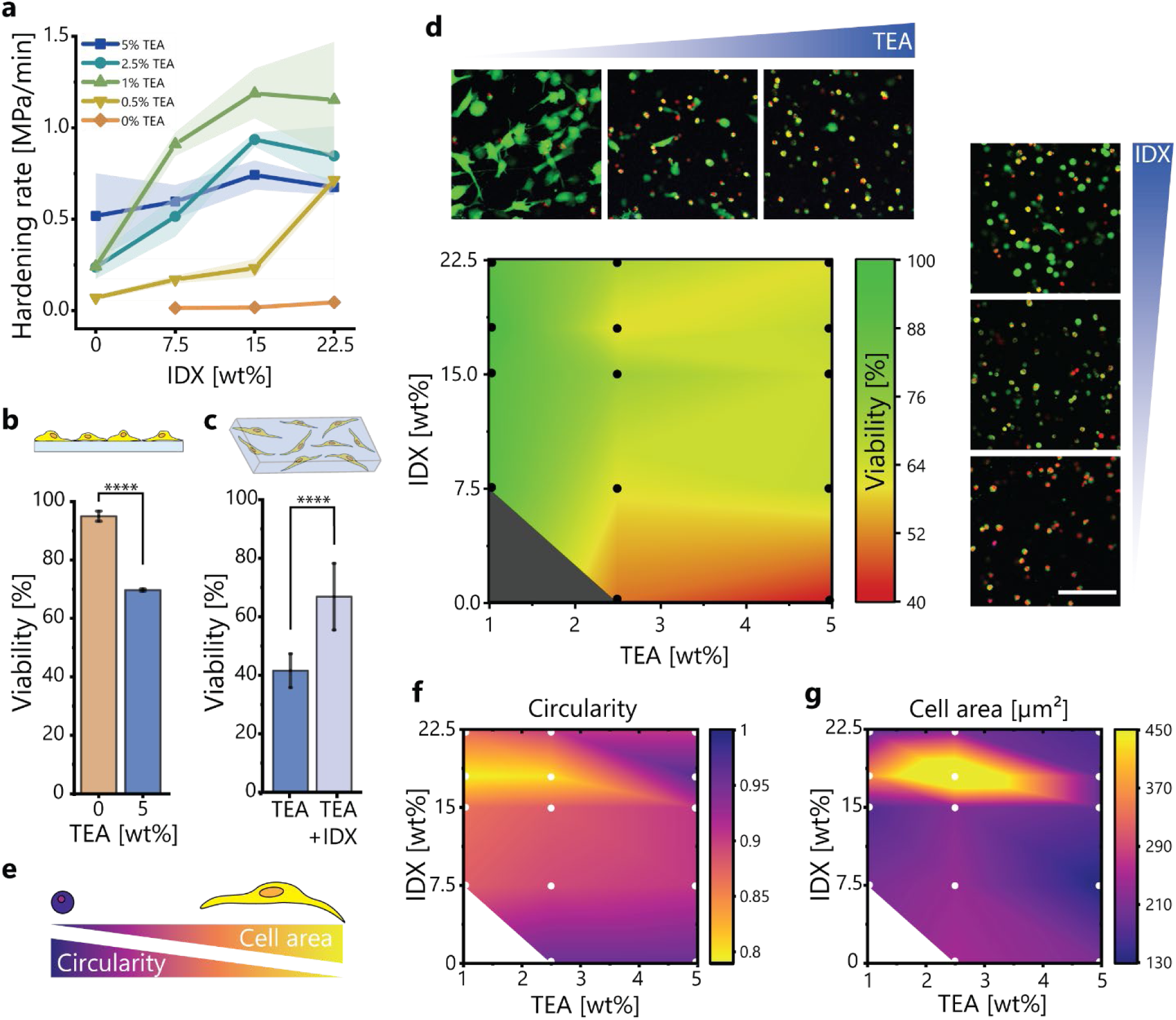
Iodixanol (IDX) improves print cytocompatibility by enabling a decrease in TEA concentration and offering an innate cytoprotective effect. (a) Dual color photorheological determination of hardening rate of various iodixanol and TEA on GelMA print formulations (n=3). (b) Viability of 3T3 fibroblast monolayer incubated with TEA (n=3). (c) Viability of 3T3 fibroblasts in 3D printed constructs containing 5 wt% TEA and optionally 18 wt% IDX. (d) Effect of iodixanol and TEA concentration on cell viability in Xolographically printed constructs. Images are live-dead (green: Calcein-AM, red: EthD-1) fluorescence micrographs of printed constructs containing (top row) 18 wt% IDX and from left to right 1 wt%, 2.5 wt%, and 5 wt% TEA and (right column) 5 wt% TEA and from top to bottom 15 wt%, 7.5 wt%, and 0 wt% IDX. (Scale bar: 50 µm) (n≥3) (e) Schematic depicting a small circular cell and a larger elongated cell. (f) Image-based analysis of the effect of iodixanol and TEA on cell circularity in printed constructs. (g) Image-based analysis of the effect of iodixanol and TEA on cell area in printed constructs. (n≥3) Data is shown as mean ± standard deviation. Analysis is via a two sample t-test. ****p<0.0001

TEA exerts substantial concentration-dependent cytotoxicity on mammalian cells (**Figure 2b**). We therefore anticipated that an iodixanol-facilitated decrease in TEA concentration would be advantageous for the cytocompatibility of Bioxolographic printing. To this end, the effects of various iodixanol and TEA concentrations on 3T3 fibroblast viability in Xolographically printed 5% GelMA hydrogels were investigated (**Figure 2d**). All conditions were printable, except for iodixanol-free samples containing 1 wt% TEA, further emphasizing the role that iodixanol plays in improving reactivity. As anticipated, viability significantly improved when decreasing TEA concentration across all tested iodixanol concentrations. Viability increased from 41% ± 6% of conventional formulations (e.g., 5 wt% TEA) to 95% ± 2% for 1% TEA and 18% iodixanol containing formulations. Interestingly, adding iodixanol aided in cell survival even when keeping TEA concentration constant (**Figure 2c**). This suggests that iodixanol possesses a previously unknown cytoprotective effect during the photocrosslinking of living materials. This is potentially mediated via antioxidant effects, preventing cells from experiencing high concentrations of free radicals^[23, 24]^. Cell shape analysis revealed that Xolographic printing with lower TEA and higher iodixanol concentrations facilitated accelerated cell spreading as observed by a reduction in cell circularity (**Figure 2e**). Similarly, printing with lower TEA and higher iodixanol concentration was associated with increased cell area (**Figure 2f**). Maximum cell area and minimum circularity were both achieved when using 18 wt% iodixanol, suggesting it as an optimal iodixanol concentration for cell-containing prints. Given the importance of cell-material interactions including cell spreading in creating tissues, this underscores the potential of iodixanol in enabling Xolographic printing of highly viable engineered living matter.

### Iodixanol enables Xolographic printing of low-photoreactivity materials

As iodixanol increases photoreactivity, we reasoned that iodixanol could also allow for the expansion of the Xolographic material toolbox by facilitating the printing of previously unprintable lower-reactivity materials. Without iodixanol, Xolography has been limited to plastic resins and three highly reactive hydrogels, namely ≥40% PEGDA, niPAAM, and ≥10% GelMA, of which only GelMA is biocompatible.^[13, 14]^ To compare the reactivity of various photopolymerizable hydrogels, dual-color photorheology was performed to determine the exposure time required to start hardening of 15% human platelet lysate-methacryloyl (hPLMA), 3% alginate-methacryloyl (AlgMA), 5% gelatin methacryloyl (GelMA), and 40% poly (ethylene glycol) diacrylate (PEGDA), which under the same irradiation conditions showed hardening after 3.95 ± 0.46, 2.19 ± 0.05, 0.37 ± 0.02, and 0.19 ± 0.04 seconds, respectively (**Figure 3a**). Similarly, quantitative analysis of the time required for these materials to reach half of their maximum storage modulus confirmed that 40% PEGDA had the highest photocrosslinking rate for followed by 5% GelMA, 3% AlgMA, and 15% hPLMA (**Figure 3b**). To investigate whether iodixanol would allow for the Xolographic printing of previously unprintable materials, these materials were printed in the presence or absence of DPI and iodixanol (**Figure 3c**). In the absence of any reactivity-boosting agent, only the highly-reactive low molecular weight PEGDA proved printable in our print formulations, confirming previous reports^[13]^. Although the addition of either DPI or iodixanol both enabled printing with 5% GelMA resins, it still did not allow printing with the formulations based on other, less reactive, biomaterials. In sharp contrast, addition of both iodixanol and DPI readily allowed printing for all tested print resins, including those based on low reactive materials such as hPLMA and AlgMA. This confirms that supplementing existing formulations with iodixanol enables Xolographic printing with previously unprintable materials. Furthermore, for the first time this allowed for the Xolographic printing of human-based bioresins, which typically associate with lower reactivity. The iodixanol-driven expansion of the Xolographic material toolbox allows for selecting materials for their (biological) function, rather than their chemical reactivity.

**Figure 3.**
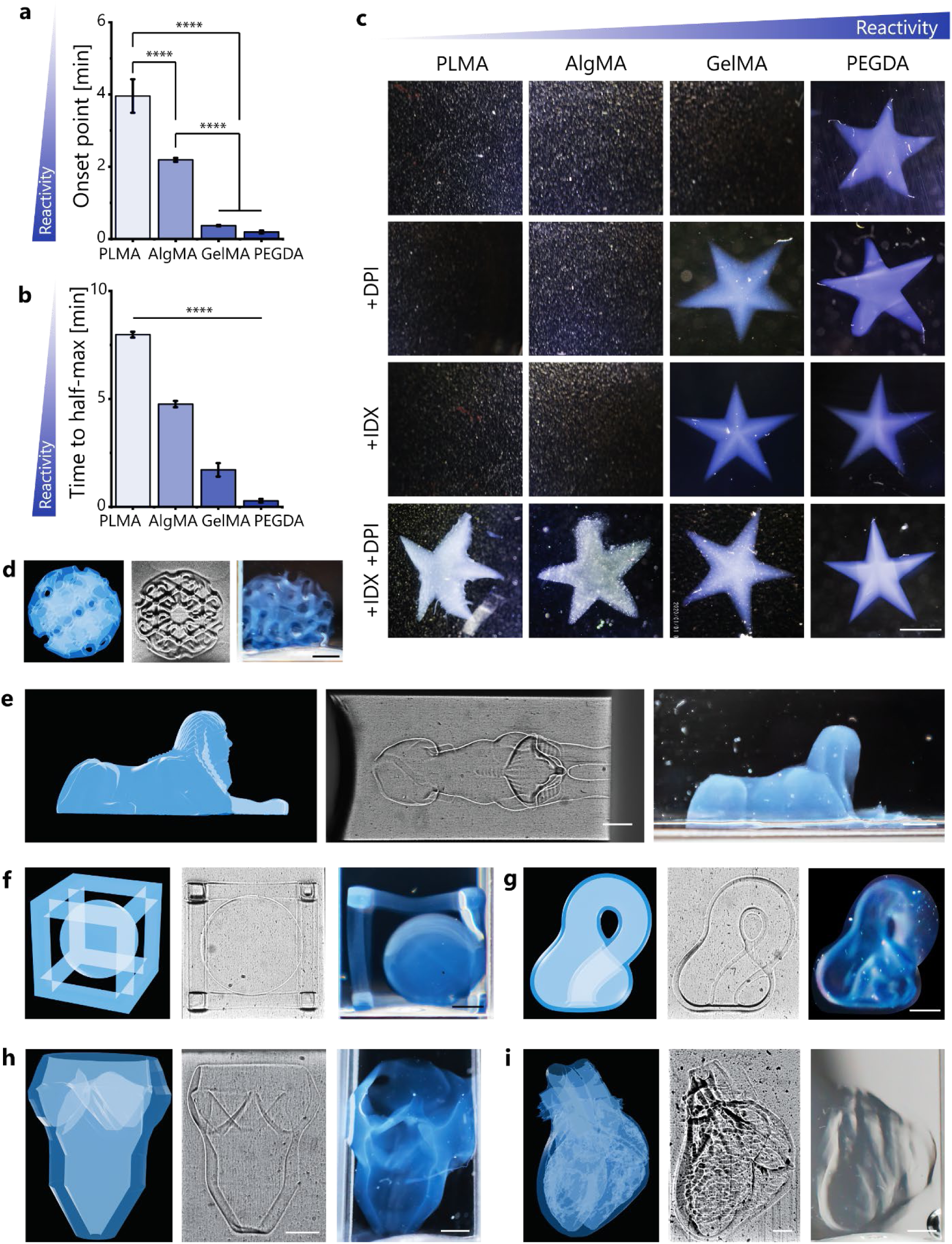
Iodixanol (IDX) enables Xolographic printing of low-reactivity materials. (a) Photopolymerization onset time for 40% PEGDA, 5% GelMA, 3% AlgMA, and 15% hPLMA. (b) The irradiation time required for each material reached half of its maximum storage modulus. (c) Example prints using print formulations based on materials of different reactivities (top row) without reactivity boosting additives, (second row) with DPI, (third row) with iodixanol, and (bottom row) with DPI and iodixanol. (d) 3D model, Schlieren photography of freshly printed constructs still submerged in their print bath, and photographs of recovered and washed structures of a gyroid, (e) sphinx, (f) a ball in a cage, (g) a Klein bottle, (h) a CT-derived tricuspid heart valve, and (i) a CT-derived healthy human heart. (Scale bars: 2 mm) Analysis is a one-way ANOVA with a post hoc Tukey test. ****p<0.0001

To demonstrate the Xolographic printability of iodixanol containing bioresins (e.g., 5% GelMA) into high resolution solid 3D objects, several unique designs including a gyroid, sphinx, tricuspid heart valve, an entire heart, a ball in a cage, and Klein bottle were printed. Schlieren photography revealed that the Xolographically printed gyroid captured both positive and negative complex features. Upon extraction from the print bath, these features were maintained (**Figure 3d**). Similarly, the sphinx revealed that both large features such as the overall design and small fine features including the striations on the headdress are captured in printing and maintained upon recovery from the print bath (**Figure 3e**). A ball in a cage showed the printing of complex, multicomponent parts, which are difficult to produce by conventional methods, such as casting. The ball and cage were printed as two separate parts and the ball floated freely within the cage (**Figure 3f**). A 3D version of a Klein bottle, a one-sided bottle without a true interior, was printed and subsequently recovered (**Figure 3g**). To show printability of biological structures, we printed a CT-derived tricuspid heart valve, with thin yet stable features, namely three individually resolved valve leaflets (**Figure 3h**). This demonstrated the ability of iodixanol containing bioresins to print high resolution and high aspect ratio designs with great accuracy and fidelity. Furthermore, we successfully printed a miniaturized version of a CT-derived heart of a healthy 17-year-old girl (**Figure 3i**). These prints accurately replicated the intricate, multichambered architecture’s positive and negative features typical for the human heart, such as two hollow ventricles and two hollow atria, including the natural variations in thickness in the heart wall. Of note, due to the hollow chambers and intricate features, this model is difficult to produce by methods other than volumetric printing. The printing of the biological structures combined with highly biomimetic materials shows the promise of Xolography for the production of tissue models. The addition of iodixanol therefore unlocks a variety of heretofore unprintable hydrogels and designs for Xolographic bioprinting.

### Iodixanol enables accurate Xolographic printing of highly cell-dense matter

Bioprinting inherently requires the use of cells in the bioresin, which poses a challenge for light-based printing techniques, owing to the light diffraction that results from refractive index mismatch between cells and the surrounding polymer solution (**Figure 4a**). The resulting light scattering lowers the print resolution in a cell concentration-dependent manner, preventing the printing of highly cell-dense living matter. This limits light-based volumetric printing techniques, including Xolography and computed axial lithography, to low cell densities compared to other biofabrication techniques.^[13, 14, 16, 20, 25]^ We hypothesize that by combining the linear nature of Xolography’s printing system with refractive index matching, we could overcome the currently unresolved cell density limitation of volumetric printing. Advantageously, iodixanol is known to reduce cell-mediated light scattering via refractive index matching^[16, 17]^. By incorporating iodixanol into the print bath, the refractive index of the print bath can be raised to match the refractive index of the cellular cytoplasm, thereby reducing light scattering. Indeed, the iodixanol increased the print resin’s refractive index from 1.348 up to 1.392 in a linear concentration-dependent manner (**Figure 4b**). Cell cytoplasm typically shows a refractive index of 1.36-1.39^[17]^, meaning that this range of iodixanol concentration allows for the matching of the refractive index of virtually all cell types. To assess how light scattering is affected by iodixanol, light transmission through bioresins containing 10 M 3T3 fibroblasts mL^-1^ was measured. (**Figure 4c**) Given high transmittance for all cell-free resins, light scattering rather than absorbance was assumed to predominate in cell-containing resins. This revealed low visible light transmittance (i.e., high scattering) for iodixanol concentrations below 15 wt% and above 22.5 wt%. Importantly, at iodixanol concentrations between 17 and 20 wt%, transmittance drastically increased, suggesting that refractive index was matched in this range. Visual inspection verified that bioresins containing this amount of iodixanol became notably less turbid and allowed for observing line patterns behind the resin, in marked contrast to all other samples (**Figure 4e**). Moreover, to confirm that the increased transparency was due to matching of refractive index between polymer and cells, holotomography was used to measure and visualize the relative refractive index of the cells in a spatially resolved manner at the subcellular level (**Figure 4f**). This revealed that while both intracellular and intercellular differences in refractive properties existed, the addition of 18 wt% iodixanol resulted in optimal refractive index matching of cells in GelMA bioresin (**Figure 4d**). Moreover, holotomography revealed that minimal refraction of the living bioresin was achieved when iodixanol was matched to the intracellular compartment, as indicated by the observation that only the area in which the cellular membrane was located remained detectable for most cells. Advantageously, this also suggested that the optical matching occurred at similar iodixanol concentrations that most effectively improved photoreactivity, cytocompatibility, and cell spreading.

**Figure 4.**
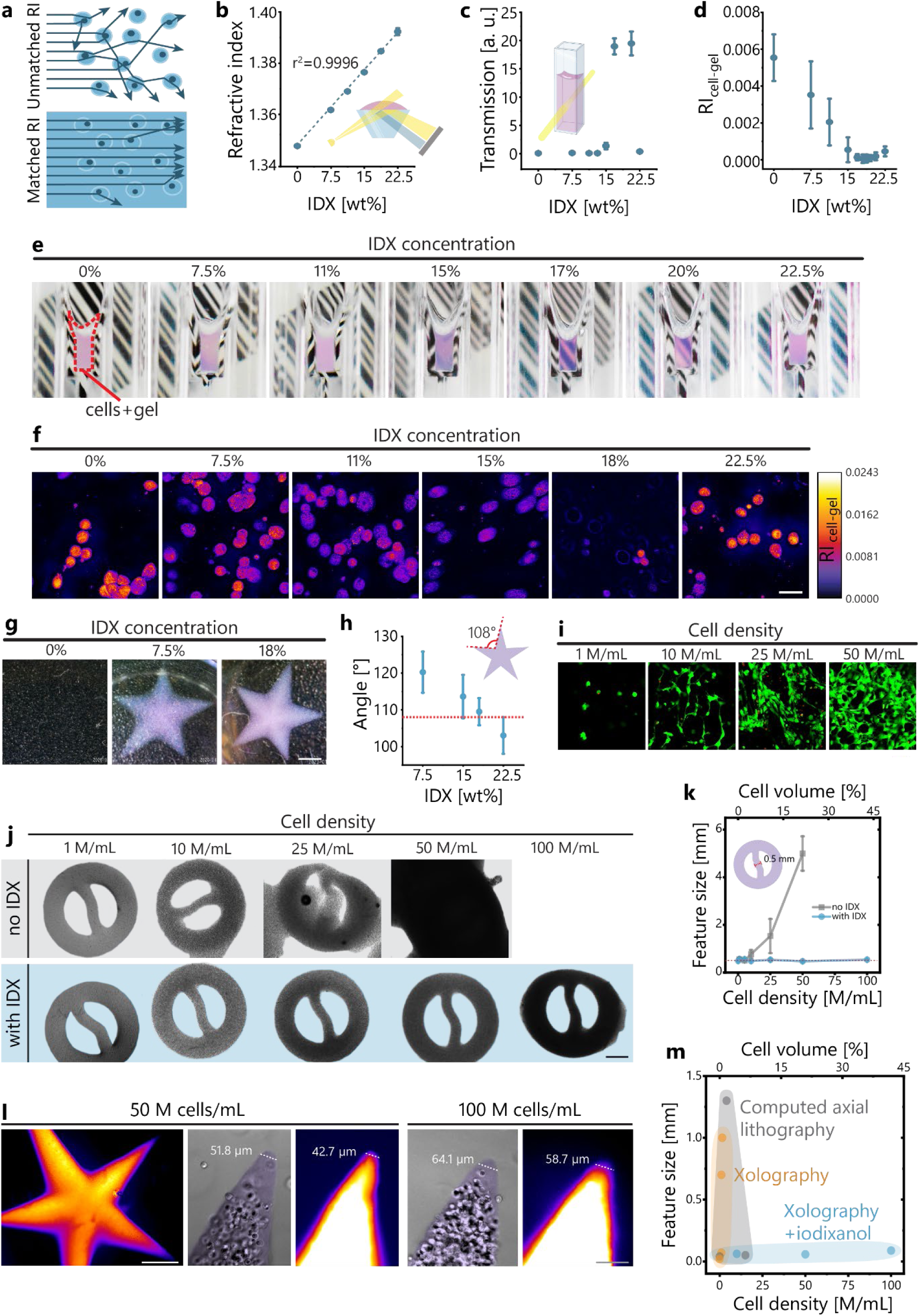
Iodixanol (IDX) reduces cell-mediated light scattering via refractive index matching, enabling accurate Xolographic printing of highly cell-dense living matter. (a) Schematic depiction of mechanism of refractive index matching. (b) Effect of iodixanol on the refractive index of the 5% GelMA bioresin (n=4). (c) Effect of iodixanol on the transmitted light through 5% GelMA bioresin containing 10 M 3T3 fibroblasts mL^-1^ (n=3, each measured four times). (d) Difference between fibroblasts refractive index and gel refractive index versus iodixanol concentration (n=C). (e) Photographs of print baths composed of 5% GelMA bioresin containing 10 M 3T3 fibroblasts mL^-1^ and various concentrations of iodixanol (IDX) in front of a barcode. (f) Holotomographic micrographs of 3T3 fibroblasts in 5% GelMA bioresins containing various amounts of iodixanol (scale bar=30 µm). (g) Xolographically printed star-shaped GelMA hydrogels containing 10 M 3T3 fibroblasts mL^-1^ and 1% TEA with 0%, 7.5%, and 18% iodixanol, where 0% iodixanol proved unprintable (scale bar: 1 mm). (h) Image-based analysis of print fidelity as measured by angle of intersection between star tips (scale bar: 1 mm). (n=5) (i) Fluorescent micrographs of live-dead stained printed constructs containing varying concentrations of 3T3 fibroblasts (scale bar: 100 µm). (j) Constructs Xolographically printed with and without iodixanol with cell densities of 1, 10, 25, 50, and 100 M 3T3 fibroblasts mL^-1^ (scale bar: 1 mm). (k) Effect of cell density on shape fidelity for both iodixanol-containing, refractive index matched resins, and iodixanol-free, non-matched resins. (l) Fluorescence and bright-field micrographs of stars Xolographically printed with 50 M and 100 M 3T3 fibroblasts mL^-1^ (scale bars: 1 mm for the whole star and 100 µm for the star tips). (m) Comparison of minimum reported feature size for cell-laden volumetric printing techniques including computed axial lithography with computational^[25]^ and iodixanol-based^[1C]^ scattering compensation, Xolography without refractive index matching^[13, 15, 2C]^, and this work employing Xolography plus iodixanol for refractive index matching.

To determine the effect of various iodixanol concentrations on print fidelity, star-shaped GelMA hydrogels containing 10 M 3T3 fibroblasts mL^-1^ were printed (**Figure 4g**). Without iodixanol, this formulation lacked sufficient reactivity to print, underscoring the role of iodixanol in boosting bioresin reactivity. To quantify shape fidelity, the exterior angle of the star was compared to the design angle (**Figure 4h**), revealing reduced sharpness and larger than ideal angles in unoptimized conditions. At optimized iodixanol values, the angle in printed constructs near-perfectly matched the ideal angle.

Although the printing of high cell densities is imperative to the biofabrication of functional living tissues, it has so far remained impossible to achieve this using Xolography due to light-scattering, which restricted printable formulations to ≤1 M cells mL^-1^ (<0.5% cells by volume) Although computed axial lithography has successfully printed 2.5-15 M cells mL^-1^ (∼1-5% cells by volume) with the use of refractive index matching and computational light-scattering compensation, it too remains far below the cell density of virtually all tissue types. Given the linearity of Xolography’s light projection, we reasoned that when effective refractive index matching is implemented, Xolography should allow for volumetric bioprinting at unprecedented (e.g., high) cell densities.

To determine how cell density affects shape fidelity with and without refractive index matching, we printed cell densities ranging from 1M to 100 M cells mL^-1^ (∼0.4-40% cells by volume) in the presence or absence of iodixanol (**Figure 4i-k**). Without iodixanol, shape fidelity drastically drops around 10 M cells mL^-1^, and no shape could be resolved at 50 M cells mL^-1^. In contrast, in the presence of iodixanol, high shape fidelity was maintained up to at least 50 M cells mL^-1^, and only minimal effects on shape fidelity were observed when printing with 100 M cells mL^-1^. Of note, 100 M cells mL^-1^ equals a cell volume fraction of >40% of the bioresin for cells that are 20 µm in diameter. This represents the (near) limit of cell-in-material printing: higher cell fractions call for alternate biofabrication strategies, such modification of cell membranes for direct cell-cell crosslinking ^[26, 27]^. To determine the print resolution at high cell densities, we printed five-point stars and quantified their tip size (**Figure 4l**). Quantification of tip size from both fluorescently labeled and bright field images revealed a minimal feature size of ∼50 µm at 50 M cells mL^-1^ (>20% cells by volume) and ∼65 µm for 100 M cells mL^-1^ (>40% cells by volume). When compared to all previously reported Bioxolographic printing approaches, this represents a two order of magnitude cell density increase while maintaining similarly high print resolution, thus opening a previously inaccessible biofabrication window (**Figure 4m**). When compared to computational or refractive-index matching scattering reduction approaches used in computed axial lithography, this represents an order of magnitude increase in cell density, without sacrificing resolution. The simple solution of combining Xolography with iodixanol-based refractive index matching thus enables bioprinting at the limit of cell-in-material printing, while maintaining sub-100 µm print resolution, enabling high resolution printing with tissue relevant cell densities.

### Iodixanol enables printing with functional hiPSC-CMs with tissue-level functions

Until now, Bioxolography has remained limited to the fabrication of GelMA hydrogels at low cell density (e.g. ∼0.5 M cells mL^-1^), which only allows for individual cellular function as it hinders cell-cell interactions. Indeed, tissue-level function of Xolographic prints has remained inaccessible due to the current inability to print highly cell dense constructs or highly transient materials (e.g., those that rapidly and dynamically remodel), both of which are essential to efficiently form cell-cell contacts with living prints. We therefore investigated whether iodixanol-enabled printing with high cell density in highly transient materials could create engineered living matter capable of tissue-level function. To this end, we selected human induced pluripotent stem cell-derived cardiomyocytes (hiPSC-CMs) as these cells demand a high cell density and fast-paced material transience to achieve tissue-level function (e.g., tissue contraction). hiPSCs were differentiated into cardiomyocytes by sequentially adapting their chemical microenvironments over time (**Figure 5a**). The hiPSCs lost expression of pluripotency markers SOX2 and OCT4 and gained expression of cardiomyocyte markers cardiac troponin T (cTnT) and striated α-actinin, indicating sarcomere formation and successful hiPSC-CM differentiation (**Figure 5b, 5c**). Moreover, flow cytometry confirmed that the resulting hiPSC-CM population was differentiated at a high level of purity, with 85% of cells expressing cTnT (**Figure 5d**).

**Figure 5.**
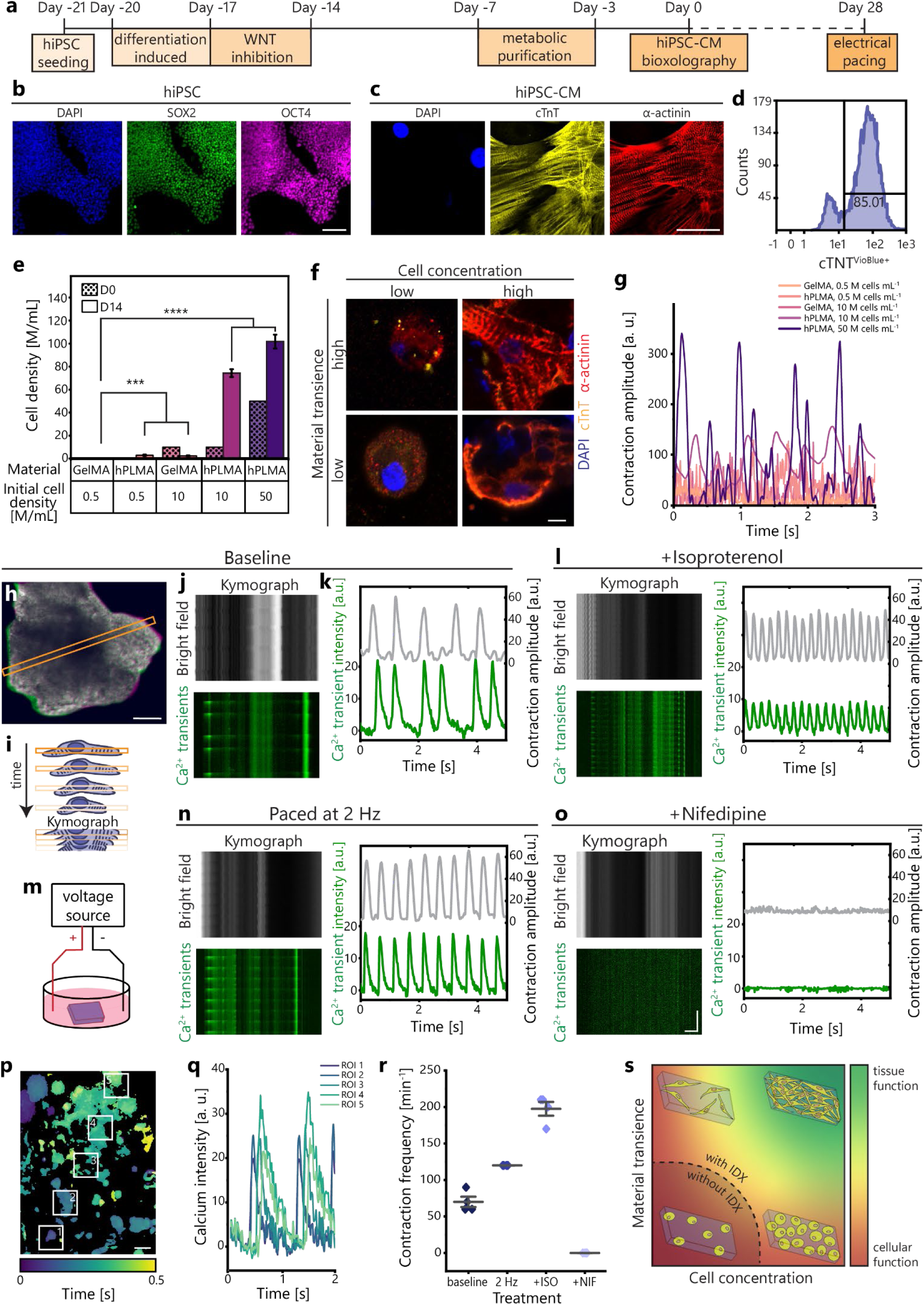
Iodixanol supplementation enables Xolographic printing of hiPSC-CMs to produce long-term stable contractile engineered living matter with tissue-level functions. (a) Schematic depicting the experimental timeline from hiPSC-CM differentiation to Bioxolography to electrical pacing of Xolographically printed engineered living matter. (b) Fluorescence micrographs of hiPSCs stained for DAPI, SOX2, and OCT4 (scale bar: 100 µm) and (c) hiPSC-CMs stained for DAPI, cTnT, and α-actinin (scale bar: 50 µm). (d) Flow cytometry histogram of hiPSC-CMs used for printing based on cardiac troponin (cTnT) expression. (e) Nuclear density of Xolographically-printed engineered living matter 14 days post-fabrication. (f) Fluorescence confocal micrographs of printed engineered living matter stained for DAPI, cTnT, and α-actinin (scale bar: 50 µm). (g) Spontaneous contraction of printed constructs after 28 days of culture as measured via MUSCLEMOTION[2S]. (h) Overlay of uncontracted (green) and contracted (magenta) printed engineered living matter (scale bar: 100 µm). (i) Schematic depicting the creation of kymograph based on captured videos. (j) Kymograph of a spontaneously contracting engineered living material with 10 M hiPSC-CMs mL^-1^ in S% hPLMA and 1% GelMA after 28 days in brightfield and with calcium transients visualized. (k) Calcium transient signal intensity and the contraction amplitude in spontaneously contracting constructs as measured by MUSCLEMOTION[2S]. (l) Resulting kymographs and calcium transient signal intensity and contraction amplitude of a engineered living material treated with 10 µM of isoproterenol. (m) Schematic of construct pacing. (n) Resulting kymographs and calcium transient signal intensity and contraction amplitude of a engineered living material when paced at 2 Hz. (o) Resulting kymographs and calcium transient signal intensity and contraction amplitude of a engineered living material treated with 10 µM of nifedipine. (horizontal scale bar: 100 µm; vertical scale bar 2 s) (p) Time-coded map of calcium transient propagation in samples with 50 M hiPSC-CMs in 9% hPLMA and 1% GelMA at day 28. (scale bar: 50 µm). (q) Calcium transient signal intensity over time in regions of interest shown in (p). (r) Contraction frequency of printed engineered living materials when contracting spontaneously, electrically paced at 2 Hz, treated with isoproterenol, or treated with nifedipine. (s) Schematic depiction of the material transience and cell density requirements to achieve tissue-level function.

To compare tissue formation in Xolographic constructs composed of high and low transience materials with high and low cell density, we printed with 0.5 M, 10 M, or 50 M hiPSC-CMs mL^-1^, representing the maximum cellular volume fraction for this cell type mixture, in low-transience 10% GelMA or a high-transience combination of 9% hPLMA and 1% GelMA. Human adult cardiac fibroblasts (haCFs) were added at 10% of the hiPSC-CM concentration to improve cardiomyocyte maturation and cell-driven material remodeling, without inhibiting cardiac conduction.^[28]^ Following Xolographic printing, hiPSC-CMs and haCFs exhibited a high level of cell viability (80.6 ± 5.7% (n=3)) (**Figure S5**). Furthermore, analyzing the nuclear density in constructs after 14 days of culture revealed that the cell concentration dropped from their initial density in GelMA-only constructs, whereas the cell concentration in hPLMA-based constructs rose, regardless of their initial cell concentration (**Figure 5e**). However, when 0.5 M cells mL^-1^ were used cell density remained low, regardless of the choice of material. With at least 10 M cells mL^-1^, cell density increased to over 70 M cells mL^-1^ when printed in hPLMA-based materials, confirming that both high cell density and transient materials are required to generate and maintain highly-cell dense constructs. Immunostaining revealed low expression of α-actinin and cTnT in prints containing 0.5 M cells mL^-1^, and high expression in prints with at least 10 M cells mL^-1^ (**Figure 5f**). Importantly, only constructs composed of highly transient material and containing high cell density formed sarcomeres, which are essential for contractility and thus indicates improved cardiomyocyte maturity. Indeed, only when printing high cell densities (≥10 M cells mL^-1^) in highly transient materials did multi-cell contractions occurred, which is a key cardiac tissue-level function (**Figure 5g**, **Movie S1**). This supports that both transient materials and high cell densities are required to achieve tissue-level function in Xolographic prints.

When a minimum cell density of 10 M hiPSC-CMs mL^-1^ were printed in highly transient material, the Xolographically biofabricated living materials contracted spontaneously at the construct level (**Figure 5h**) and, in regions of high cell density, synchronously at an average rate of 70±14 beats per minute at day 28, shown via kymographs and intensity-based analysis of videos of contracting constructs (**Figure 5h-k**, **Movie S1**). Moreover, calcium transient imaging revealed that the contraction occurred in a calcium-mediated manner, as expected for hiPSC-CMs (**Figure 5j**, **Movie S2**). Upon electrical pacing, the engineered living materials exhibited synchronous contractions and calcium transients that intimately followed the pacing rate (**Figure 5m,n**, **Movie S3**). To assess whether constructs tissue-level function was responsive to pharmacological stimulation, engineered living materials were treated with 10 µM of β-adrenergic agonist isoproterenol (**Figure 5l, Movie S3**), and subsequently with 10 µM of L-type calcium channel blocker nifedipine (**Figure 5o, Movie S3**). When treated with isoproterenol, contraction frequency increased to 198±19 beats per minute. Upon subsequent treatment with nifedipine, contractions ceased (**Figure 5r**), demonstrating expected responses to both drugs, indicating that these engineered living materials possess functional responses to pharmacological stimulation. While 10 M hiPSC-CMs mL^-1^ proved adequate for achieving regions of synchronous contraction, which follow electrical pacing and respond to pharmacological stimulation, samples with 50 M hiPSC-CMs mL^-1^ showed improved synchronicity across the sample (**Figure 5p-q**, **Movie S2**). Mapping of calcium transients over time in these samples with ultra-high cell density confirmed wave-motion contractions through the prints, confirming tissue-level function. In contrast, this behavior could not be achieved when using either low cell density or low-transience materials. Taken together, iodixanol enabled the use of highly transient materials and high cell densities, which for the first time enabled tissue-level function in Xolographically printed engineered living matter (**Figure 5s**).

## Conclusion

This study demonstrates that iodixanol boosts the reactivity of type II photoreaction systems, which allows for shorter crosslinking times and lowered coinitiator concentrations. This finding is particularly relevant for Bioxolography, where the sole addition of iodixanol improves the photoreactivity of DCPIs as well as the cytocompatibility and transparency of bioresins, enabling high resolution printing with unprecedentedly high cell densities. Improved photoreactivity allowed for Xolographic printing at significantly lower UV light doses and higher printing speeds, reducing cell exposure to UV light, cytotoxic bioresin components, and time spent outside ideal culture conditions. Additionally, the improved reactivity allows for the printing of previously unprintable low-reactivity materials such as naturally-derived hydrogels including hPLMA and AlgMA. This expansion of the material toolbox unlocks further possibilities in designing and selecting materials for the desired (biological) function, rather than based on their high reactivity. The addition of iodixanol facilitated improved cytocompatibility both in an indirect manner by enabling the reduction of the required cytotoxic coinitiator, and in a direct manner, potentially by an antioxidant effect. Regardless, improved cytocompatibility correlated with enhanced cell spreading and establishment of cell-cell contacts, which is essential for the creation of engineered living tissues. Moreover, this also enabled the Xolographic printing of sensitive cells including hiPSC-derived cells, while retaining their viability, sensitivity to microenvironmental cues, and biological functionality. Furthermore, the addition of iodixanol enabled the high-fidelity and high-resolution printing of bioresins containing cell concentrations that were two orders of magnitude greater than previous possible (i.e. from 0.5-1 (<0.5% cells by volume) to 100 M cells mL^-1^ (>40% cells by volume)) via both improved photoreactivity and refractive index matching. Of note, while other volumetric printing methods such as computed axial lithography are able to achieve printing at maximal cell densities of 2.5-15 M mL^-1^ (∼0.5%-5% cells by volume), even in the presence of light scattering reduction^[16, 25]^, here we are able to print a near order of magnitude higher cell density, approaching the limit for cell-in-material printing^[26]^, enabling high resolution printing with tissue-relevant cell densities. Additionally, improved photoreactivity broadened the Xolographic material toolbox, enabling for the first time printing with highly transient materials. Taken together, the simple addition of iodixanol to bioresins allows for the biofabrication of functional highly cell-dense engineered living matter, with tissue-level, rather than individual cell-level, functions, as demonstrated by the successful printing of contractile hiPSC-based cardiac-like engineered living matter. While these findings are of particular relevance to the tissue engineering field, we anticipate that the highly water-soluble, iso-osmotic, and non-ionic nature of iodixanol as a photoreactivity boosting agent is likely to also benefit other fields using aqueous materials, including soft robotics, photovoltaics, dentistry, and wearable sensors, by offering a novel solution to improve the rate and efficacy of their type II photoinitiation.

## Materials and methods

### Cell culture

#### Fibroblasts

3T3 fibroblasts were cultured at 37 °C and 5% CO_2_ in DMEM (Gibco) supplemented with 10% FBS (Sigma-Aldrich), 100 U mL^-1^ penicillin/streptomycin (Gibco), 1 mM sodium pyruvate (Sigma-Aldrich), and 0.14% 2-mercapto-ethanol (Gibco). At 80% confluency, cells were harvested with 0.25% Trypsin-EDTA (Gibco).

#### Cardiomyocytes differentiation and culture

Human induced pluripotent stem cells (hiPSCs; Coriell, GM25256) were maintained in Essential 8 (E8) medium (Thermo Fisher, A1517001) on vitronectin (1:100 in PBS; Thermo Fisher, A31804)-coated 6-well plates. Differentiation to hiPSC-derived cardiomyocytes (hiPSC-CMs) was induced as described previously.^[30]^ Briefly, hiPSCs were seeded at a density of 15,000–25,000 cells per cm^2^ on Matrigel (1:100; Corning, 354230) coated 6-well plates in E8 medium. After 24 h (day -20), mesodermal differentiation was induced by Activin-A (20-30 ng mL^-1^, Miltenyi, 130–115-010), BMP4 (20 ng mL^-1^, RCD systems, 314-BP/CF), and WNT activator CHIR99021 (1.5-2.25 μmol L^-1^, Axon Medchem, 1386) in BPEL medium^[31]^. At day -17, cells were refreshed with BPEL containing WNT inhibitor XAV939 (5 μmol L^-1^, RCD Systems 3748). Cells were refreshed with BPEL on day -14 and -10 of the differentiation. From days -7 to -3, the differentiated cells were metabolically purified in previously defined cardiomyocyte medium^[30]^ supplemented with 5 mM sodium DL-lactate solution (60%, Sigma-Aldrich, L4263). From days -3 to 0 and post printing, the hiPSC-CMs were maintained in cardiomyocyte medium supplemented with lactate and 4.5 mM glucose. The cells were then dissociated with TrypLE 10X (ThermoFisher, A1217702) and then used for printing. Cells were reserved for analysis by flow cytometry to ensure >80% positive expression of cardiac troponin T (cTnT).

#### Human cardiac fibroblasts

haCFs (Promocell, C-12375) were maintained in Fibroblast Growth Medium-3 (PromoCell, C-23025). Medium was refreshed every 48 hours until 70-90% confluency was reached, when they were then dissociated with TrypLE 10x. The cells were passaged and expanded until reaching 11 passages, prior to cryopreservation. Just prior to use in the printing process, haCFs were thawed.

### Solubility

Solubility of iodixanol (Sigma-Aldrich) and diphenyliodonium chloride (DPI; Sigma-Aldrich) in water was determined by sonicating a mixture of water and solid DPI or iodixanol for 30 minutes. The solution was then centrifuged, and the supernatant was removed. The supernatant was then lyophilized overnight to determine mass of the dissolved solids.

### Alginate modification

Alginate methacrylation was conducted as previously described.^[32]^ Briefly, sodium alginate (2 g) (Alfa Aesar, A18565) was dissolved at 1% in 50 mM MES + 0.5M NaCl buffer, followed by 1-ethyl-3-(3-dimethylaminopropyl)-carbodiimide hydrochloride (EDC; Sigma-Aldrich) and *N*-hydroxysuccinimide (NHS; Sigma-Aldrich). After 5 minutes, 2-aminoethyl methacrylate (AEMA; Sigma-Aldrich) (molar ratio 1:2:1 NHS:EDC:AEMA) was added and the reaction allowed to proceed. After 24 h, the reaction mixture was precipitated in acetone, then desiccated overnight. The desiccated product was dissolved in deionized water and dialyzed against deionized water for 3 days (MWCO 1 kDa), filtered at 0.22 µm, then lyophilized. The alginate methacrylol (AlgMA) was then dissolved in ultrapure water for a final concentration of 3 wt% in further experiments.

### Xolography

Gelatin methacryloyl (GelMA; 167 kDa, 86% degree of functionalization, Rousselot X-Pure) was dissolved in PBS at 37 °C. DPI was dissolved in dimethylsulfoxide (DMSO; Hybri-Max, Sigma-Aldrich) under stirring at 60 °C for 1 h. Dual color photoinitiator (DCPI; DCPI 5002, xolo GmbH) was dissolved in triethanol amine (TEA; Sigma-Aldrich) for 1 h under stirring. Iodixanol (Optiprep, Stemcell), *N*-vinyl-2-pyrrolidinone (VP; Sigma-Aldrich), and TEA were used as commercial solutions. Stock solutions were then mixed to make the formulations as described. Unless otherwise mentioned, print formulations used 5 wt% GelMA, 0.015 wt% DCPI, 1 mM DPI, and 0.3 wt% VP, with variable amounts of TEA and iodixanol. When used, alginate methacrylate was dissolved in ultrapure water, and used at a final concentration of 3 wt%. After printing, recovered AlgMA samples were post-crosslinked with CaCl_2_. Human platelet lysate methacryloyl (hPLMA; Metatissue) was dissolved at room temperature in PBS and incorporated at a final concentration of 15 wt%. Poly(ethylene glycol) diacrylate (PEGDA; M_n_ 575 g mol^-1^, Sigma-Aldrich) was used at a concentration of 40 wt%.

Prepared formulations were then loaded into UV-Vis transparent cuvettes (Brand) and refrigerated for 30 minutes to ensure GelMA gelation, and allowed to reach room temperature for a minimum of 30 minutes prior to Xolographic printing. Printing was performed using a Xube (xolo GmbH) Xolographic printer. Prints were recovered by washing repeatedly (>3x) with PBS at 37 °C to remove uncrosslinked resin.

The model used for the sphinx print was created by Thingiverse user cerberus333 under Creative Commons - Attribution - Non-Commercial.^[33]^ The trileaflet valve was modified from Lee et al.^[34]^ The human heart model was used from NIH 3D repository.^[35]^

#### Print characterization

To determine print quality, recovered prints were imaged using a digital microscope (Andonstar AD409) or a digital camera (Canon EOS RP) equipped with a RF 24-105mm F4-7.1 IS STM-lens. To determine print fidelity of cell-containing stars, the exterior angle of the stars were measured from tip to the meeting point of two tips. Circular constructs with interior struts were imaged with an inverted microscope (EXI-310, ACCU-SCOPE). High cell density stars were stained post swelling with NHS ester Alexa Fluor 647 (Lumiprobe) and imaged using a fluorescence microscope (EVOS FL Imaging system microscope, Thermo Fisher). Images were adjusted using Lightroom and Photoshop (Adobe) to improve visualization.

### Photorheology

The reactivity of print formulations was measured via rheology during irradiation by UV and visible light. Measurements were performed using a Discovery HR20 rheometer (TA Instruments) equipped with 12 mm parallel plates, the top one stainless steel, the bottom quartz. The formulation was pipetted onto the plate and allowed to reach room temperature. Time sweep measurements were performed at an oscillation strain of 1% and frequency of 1 hz. For DCPI-containing samples, one minute into the time sweep, the samples were irradiated with UV and visible light from a 6-wavelength high-power LED source (Chrolis, Thorlabs). UV light at 365 nm and 37 mW cm^-2^ was pulsed on for 3 seconds, then off for 30 seconds for the duration of the measurement. Visible light at 565 nm with an intensity of 140 mW cm^-2^ was on for the duration of the measurement. Hardening rate was determined from slope of the region of linear increase in storage modulus.

To assess the effect of iodixanol on other photoinitiators, formulations were prepared with varying amounts of iodixanol. A formulation containing 0.25 wt% lithium phenyl-2,4,6-trimethylbenzoylphosphinate (LAP; Sigma-Aldrich) in 25 wt% PEGDA and 0-15 wt% iodixanol was exposed to constant light at 365 nm and 37 mW cm^-2^. For eosin Y, a formulation of 2.5 µM eosin Y (Sigma-Aldrich) and 2.5 wt% TEA, 10 wt% PEGDA, and 0-15 wt% iodixanol was exposed to constant visible light at 565 nm and 21 mW cm^-2^. For DCPI and BisTris (2-[bis(2-hydroxyethyl)amino]-2-(hydroxymethyl)propane-1,3-diol, Thermo Fisher), TEA was replaced with 8.4 wt% BisTris. For methylene blue (Sigma-Aldrich) and TEA, 20 wt % PEGDA was crosslinked with 0.01 wt% methylene blue, 3% TEA, and 0-15 wt% iodixanol and exposure to to constant 625 nm light. For riboflavin, 20 wt% PEGDA was crosslinked with 9.35 mg mL^-1^ riboflavin with either 3 wt% TEA or 3.57 mg mL^-1^ sodium persulfate (SPS, Advanced BioMatrix) supplemented with either 0-15% iodixanol or 0-1000 µM DPI and exposed to constant visible light at 475 nm and 3 mW cm^-2^.

To determine material reactivity, 0.25 wt% LAP was used instead of DCPI, DPI, and VP. 15% hPLMA, 3 wt% AlgMA, 5 wt% GelMA, and 40 wt% PEGDA were used to determine reactivity, as these were the concentrations used for printability testing. Visible light at 405 nm and an intensity of 4 mW cm^-2^ was irradiated on the samples starting 1 minutes into measurements. The hardening onset point was defined as the point at which the rheological curve deviated from its initial baseline behavior. This was determined using TRIOS software by fitting two lines and determining the time at which the two lines intersected. 0 s was defined as when irradiation began. The time at half maximum was the time at which the storage modulus reached halfway between its initial value and its maximum. These values were used in lieu of hardening speed here, since they give better comparisons between materials, due to independence from inherent mechanical properties.

### Bioxolography

For cell-containing prints, cell suspension was mixed with the prepared solution to yield a final cell concentration of 10 M mL^-1^, unless otherwise specified. The cell-containing print formulation was then loaded into UV-vis transparent cuvettes or microcuvettes (Brand) and briefly refrigerated to allow for hydrogel gelation, to prevent print sinkage prior to Xolographic printing as described above. Unless otherwise stated, 1 mm thick flat sheets were printed in microcuvettes. After printing, samples were recovered by immediate washing with plain DMEM at 37 °C a minimum of three times. 3T3 fibroblasts were printed in 5 wt% GelMA formulations described above. For hiPSC-CM containing prints, print formulation contained 1 wt% GelMA and 9 wt% hPLMA, 0.06 wt% DCPI 5004, 8.4 wt% BisTris, 0.3 wt% VP, and 37 wt% iodixanol. HiPSC-CMs were printed at concentrations of 0.5, 10, and 50 M mL^-1^; haCFs were printed a concentration of 10% of the hiPSC-CM concentration (i.e., 0.05, 1, and 5 M mL^-1^). Unless specified otherwise, cell-containing prints contained 1 wt% TEA, except for those shown in **Figure 4j**, which contained 2.5% TEA and 0.002 wt% 4-Hydroxy-TEMPO (Sigma-Aldrich) to ensure proper reactivity, also in absence of iodixanol.

#### Cell viability

Samples were incubated with 1 µM calcein-AM (Sigma-Aldrich) and 3 µM ethidium homodimer-1 (Sigma-Aldrich) in PBS for 30 minutes, and then washed with PBS. Printed samples were then imaged using a confocal laser scanning microscope (Zeiss 880 LSM) with a 10x Plan-APOCHROMAT objective. Cell monolayers were imaged using a fluorescence microscope (EVOS FL Imaging system microscope, Thermo Fisher). Cell viability was determined by using the Find Maxima tool in ImageJ. All samples were measured in triplicate.

To determine cell circularity and area, calcein-AM images were segmented using a user-trained Cellpose 2.0 model^[36]^. Circularity and area were then measured from the Cellpose-generated ROIs using ImageJ.

To determine cytotoxicity of printing components, 3T3 fibroblasts were incubated with the desired component dissolved in PBS for 30 minutes, then washed twice with PBS, and new medium added. 24 h later, the samples were stained for viability, as described above. Controls were incubated with PBS and washed, as described above.

#### Immunoffuorescence imaging

##### Cell monolayer

hiPSC pluripotency was confirmed using SOX2 and OCT4 staining. Successful CM differentiation was confirmed using a-actinin and cTnT. hiPSCs were fixated in 4% paraformaldehyde for 30 minutes at room temperature. Membranes were permeabilized with 0.1% Triton-X 100 (Sigma-Aldrich), washed, then blocked using 5% FBS for 2 h. Cells were incubated with anti-SOX2 Alexa Fluor 488 (1:200, Thermo Fisher, ab2574478) and anti-OCT4 (1:200, Santa Crus Biotechnology, sc-5279) overnight at 4 °C, then washed three times with PBS. hiPSCs were then incubated with secondary antibody goat-anti mouse Alexa Fluor 647 (1:500, Thermo Fisher, ab2535804) for 1 h, and then washed three times. Nuclei were then stained using DAPI (1:2500, Thermo Fisher). hiPSCs were then imaged using a fluorescence microscope (EVOS FL Imaging system microscope, Thermo Fisher). hiPSC-CMs were fixated in 4% paraformaldehyde in PBS for 20 minutes. The cells were permeated with 0.1% Triton-X 100 (Sigma-Aldrich) in PBS for 8 min and blocked with 1 vol% bovine serum albumin (BSA; Sigma-Aldrich) in PBS for 1 h. They were next incubated overnight at 4 °C with primary antibody anti-alpha-actinin (1:800; Sigma, A7811) and anti-Cardiac Troponin I (1:200; Abcam, ab10231). They were washed three times in PBS and incubated with secondary antibody Goat-anti-Mouse IgG Alexa Fluor 647 (1:500; Invitrogen, A21235) and Goat-anti Rabbit IgG Alexa Fluor 488 (1:500; Invitrogen, A27034) for 1 h at room temperature. The cells were washed three times in PBS and stained with DAPI (1:3000) for 30 min at room temperature. CMs were imaged using a confocal laser scanning microscope (Zeiss 880 LSM) with a 20x Plan-APOCHROMAT objective.

#### Printed tissue

Printed samples were fixated in 4% paraformaldehyde for 2 h at room temperature. Cells were permeated with 0.25% Triton-X 100 for 30 minutes and then blocked with 5 wt% BSA and 0.1% Tween 20 (Sigma-Aldrich) for 1 h. Next, samples were incubated with primary antibody anti-alpha-actinin (1:800; Sigma, A7811) and anti-Cardiac Troponin I (1:200; Abcam, ab10231) at 4 °C for 48 h. Samples were washed three times with PBS and subsequently incubated with secondary antibodies goat-anti-rabbit IgG Alexa Fluor 488 (1:500; Invitrogen, A27034) and goat-anti-mouse IgG Alexa Fluor 647 (1:500; Invitrogen, A21235) at room temperature for 2 h. Samples were then washed thrice with PBS and nuclei stained with DAPI (1 µg/mL) at room temperature for 30 minutes. Samples were mounted then imaged using a confocal laser scanning microscope (Zeiss LSM 880) with a 63x C-APOCHROMAT water immersion objective.

#### Electrical pacing and drug testing

28 days after printing, samples were electrically paced at a frequency of 2 Hz (10 ms biphasic pulses at 25 V cm^-1^) using two platinum wire electrodes connected to a custom-made voltage source and immersed in the culture medium. Samples were subsequently incubated with 10 µM of isoproterenol (Sigma, I5627) for five minutes, and then imaged. Then samples were incubated with 10 µM of nifedipine (Sigma, N7634) for 5 minutes, and then imaged. For visualization of calcium propagation, the tissues were incubated in PBS containing 5 µM calcium dye Fluo-8 (Abcam, AB142773), 1% (vol/vol) pluronic (Sigma-Aldrich P2443), and 15 mM glucose at 37 °C for 45 minutes. Tissues were washed once in PBS before imaging. Videos of contracting samples were acquired using a widefield fluorescence microscope (Nikon Ti2 Eclipse).

### Optical Properties

#### Refractometry

For determination of refractive index, DCPI and DPI were excluded to prevent premature sample curing and were replaced by TEA and DMSO, respectively. Prepared samples were pipetted onto the refractometer (Mettler Toledo Refracto 30GS) measurement cell. After allowing samples to reach room temperature, the refractive index was measured.

#### Holotomography

Samples were prepared as previously described. DCPI and DPI were replaced by TEA and DMSO, respectively, to prevent premature crosslinking. Cell suspension was mixed with the prepared solution and then loaded into a TomoDish (Tomocube). Relative refractive index was then imaged via holotomography (Tomocube). Cell refractive index was then measured by using ImageJ’s “Analyze Particles” function to outline cells. The refractive index in the region defined by analyze particles was then measured. Gel refractive index was measured manually in ImageJ in regions without cells.

#### Transmission

Samples were loaded into UV-vis transparent microcuvettes (Brand). Samples were loaded into a homemade transmission setup, where they were illuminated with a collimated visible light source. Transmitted light was then measured and averaged over wavelengths from 450 nm to 1000 nm.

### Statistical analysis

Data is reported as mean ± SD. Statistical significance was determined in OriginLab software with a one-way ANOVA test with an post hoc Tukey test, unless otherwise noted. Significance was defined by a value of p<0.05.

## Acknowledgements

The authors wish to thank Tom Knop for assistance with measuring bioresin transmittance, Minh Luu for confirming hiPSC pluripotency, and Danique Snippert and Anne Braam for differentiating hiPSCs to cardiomyocytes. J.L., R.P., and M.C.J. acknowledge financial support from the European Innovation Council (ELM project 210803707: BioRobot-MiniHeart). R.P. acknowledges financial support from the European Research Council (ERC, Advanced Grant Heart2Beat, project number 101098372). J.L. acknowledges financial support from the European Research Council (ERC, Advanced Grant, project number 101266635, VitaliTE).

## Notes

### Competing Interest Statement

N. F. K. is an employee of xolo GmbH. J. F. M. is a co-founder of Metatissue. C. A. C. is a co-founder and CEO of Metatissue.

