## Supporting Information for "Xolographic printing of cell-dense engineered living matter via iodixanol-driven enhancement of type II photoinitiation"

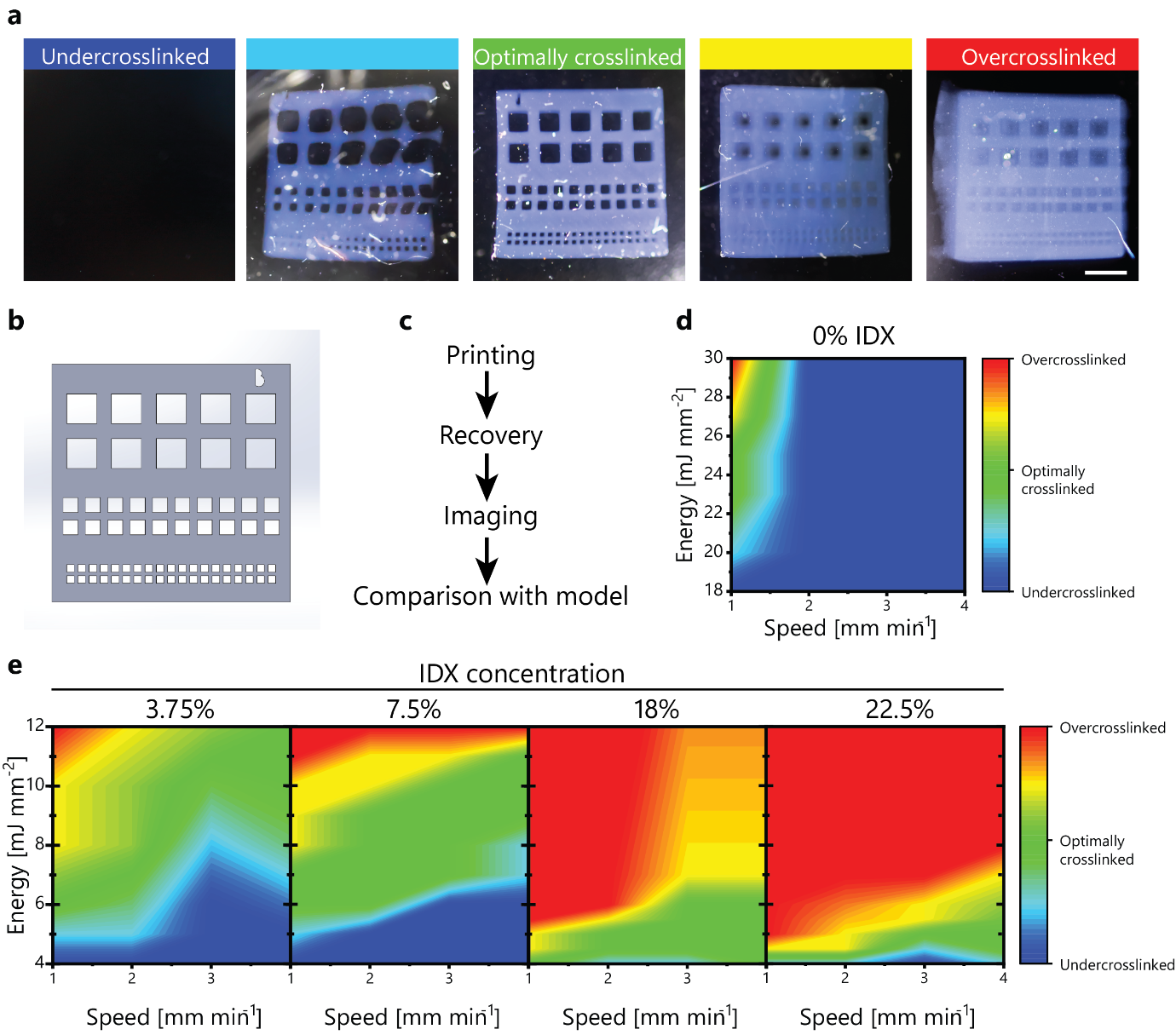

***Figure S1****. (a) Constructs used to determine printability of formulations with various amounts of iodixanol. Best quality prints are those that best resemble “optimally crosslinked” prints. (b) CAD model used to assess printability. (c) Printability assessment workflow. (d) Iodixanol-free print map. (e) Print maps for GelMA-based formulations containing varying amounts of iodixanol.*

### Photorheology

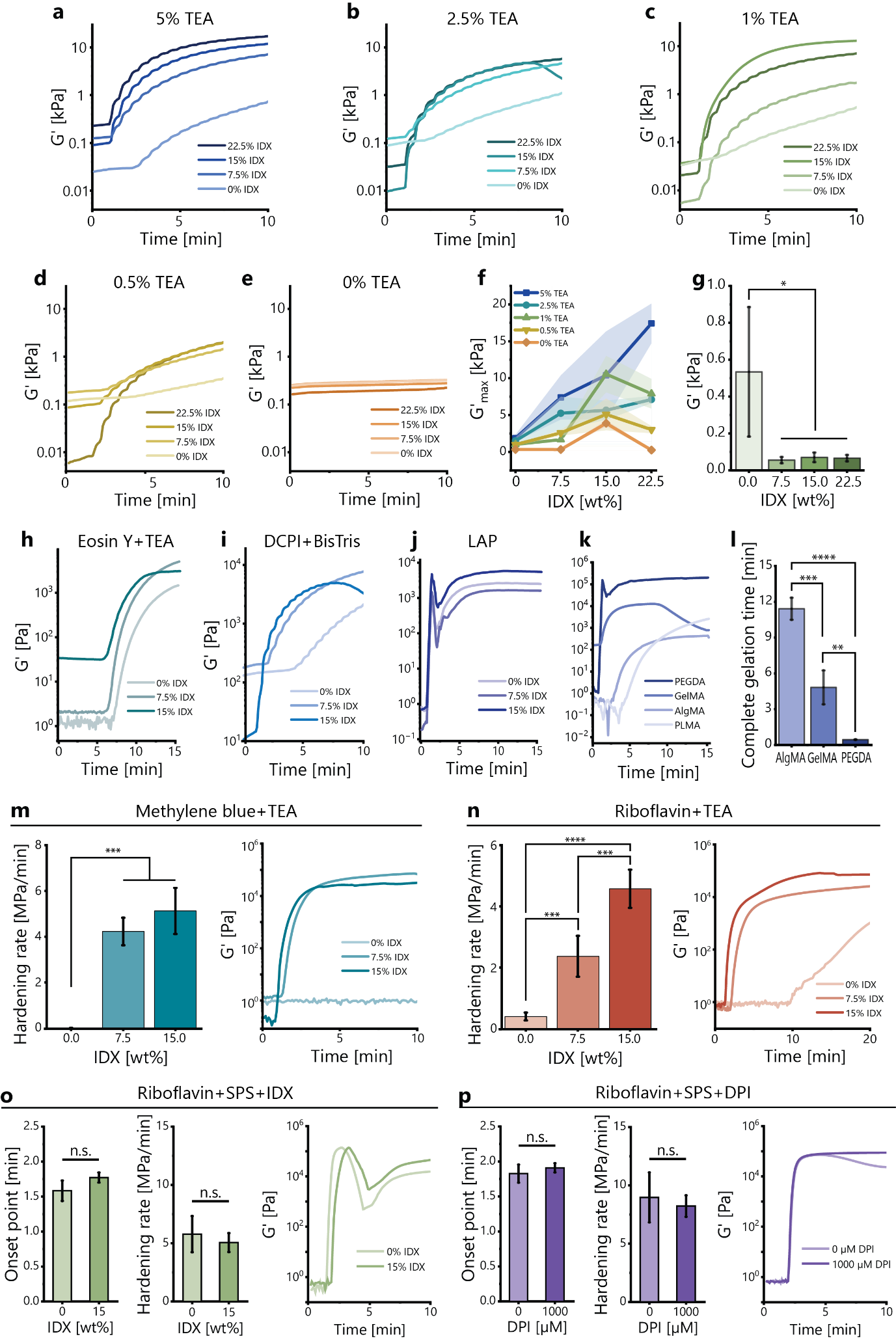

***Figure S2.*** *Dual color photorheology of GelMA-based print formulations with varying amounts of iodixanol and (a) 5% TEA, (b) 2.5% TEA, (c) 1% TEA, (d) 0.5% TEA, or (e) 0% TEA. (f) Final stiffness of GelMA-based print formulations for various iodixanol and TEA concentrations using dual color photorheology (n=3). (g) Influence of iodixanol on the storage modulus of printed and recovered constructs (n=3). Effect of iodixanol on the hardening behavior of (h) 10% PEGDA with eosin Y and TEA, (i) 5% GelMA with DCPI and BisTris, or (j) 10% PEGDA with LAP. (k) Photorheological behavior of 40% PEGDA, 5% GelMA, 3% AlgMA, and 15% hPLMA. (l) The required time for each material to gel completely, based on the first derivative of the storage modulus (n=3). The completed gelation time of hPLMA exceeded the experimental duration of 15 minutes. (m) Effect of iodixanol on the hardening rate of 20% PEGDA using methylene blue and TEA and (n) riboflavin and TEA for photoinitiation. (o) Effect of iodixanol and (p) DPI on the hardening behavior of 20% PEGDA using riboflavin and SPS as a photoinitiation system. Analysis is a one-way ANOVA with a post-hoc Tukey test. *p<0.05, **p<0.01, ***p<0.001, ****p<0.0001*

### UV-Vis analysis of iodixanol reactivity enhancement

#### To investigate the mechanisms by which iodixanol increases the reactivity of type II photoinitiation systems, UV-vis spectroscopy was performed under concurrent irradiation with visible light. Irradiation with visible light gives insight into the mechanism after the DCPI transitions to the merocyanine phase (**Figure S3a**). In aqueous systems, DCPI 5002 exists in both the spiropyran and merocyanine forms (**Figure S3b**), due to thermally-driven transition to the merocyanine phase, enabling insight into the mechanism with a single visible light source. Conventional formulations using DCPI and BisTris as a photoinitiation system start polymerizing after 1200 s, while polymerization starts around 200 s when the formulation is supplemented with iodixanol (**Figure S3c-e**). Without BisTris to function as a coinitiator, iodixanol and DCPI do not cause polymerization (**Figure S3f**). This indicates that iodixanol does not act as a coinitiator, but improves the rate of polymerization due to another effect, such as via a synergistic reaction with the BisTris or by oxidizing the DCPI back to a state where it can participate in polymerization.

#### Methods

To assess the effect of iodixanol on the polymerization process, UV-vis spectroscopy (Cary 50, Varian) was performed under perpendicular illumination at 565 nm (LED; Thorlabs), collimated with an adjustable collimation adapter with an AR-coated lens (Thorlabs, SM2F32-A). Power output was maintained at constant values via a T-Cube LED driver (LEDD1B, Thorlabs). Stirring was maintained at 1200 rpm and temperature was maintained at 25 °C via the cuvette holder (Luma 4, Quantum Northwest). To determine the effect of iodixanol on the dual-color DCPI-driven polymerization process while excluding any effects of IDX in the UV range, spectroscopy was performed during perpendicular illumination at 565 nm. At 25 °C, DCPI 5002 exists in both the dormant spiropyran form and in the active merocyanine form (Figure S2b), and therefore irradiation with visible light alone can initiate polymerization. To assess polymerization under visible light, formulations of 4.5% poly(ethylene glycol) diacrylate (PEGDA; Sigma-Aldrich), 5.5% acrylamide/bisacrylamide (Sigma-Aldrich), and optionally 364 mM 2-bis(2-hydroxyethyl)amino-2-(hydroxymethyl)-1,3-propanediol (BisTris; TCI Deutschland GmbH), 0.087% dual-color photoinitiator (DCPI 5002; xolo GmbH), and/or 21.8% iodixanol (Proteogenix S.A.S.) were loaded into fluorescence cuvettes (Brand). To assess polymerization, absorbance between 700 and 749 nm (above the range of DCPI and IDX absorbance) was integrated and taken to represent scattering due to PEGDA/acrylamide polymerization.

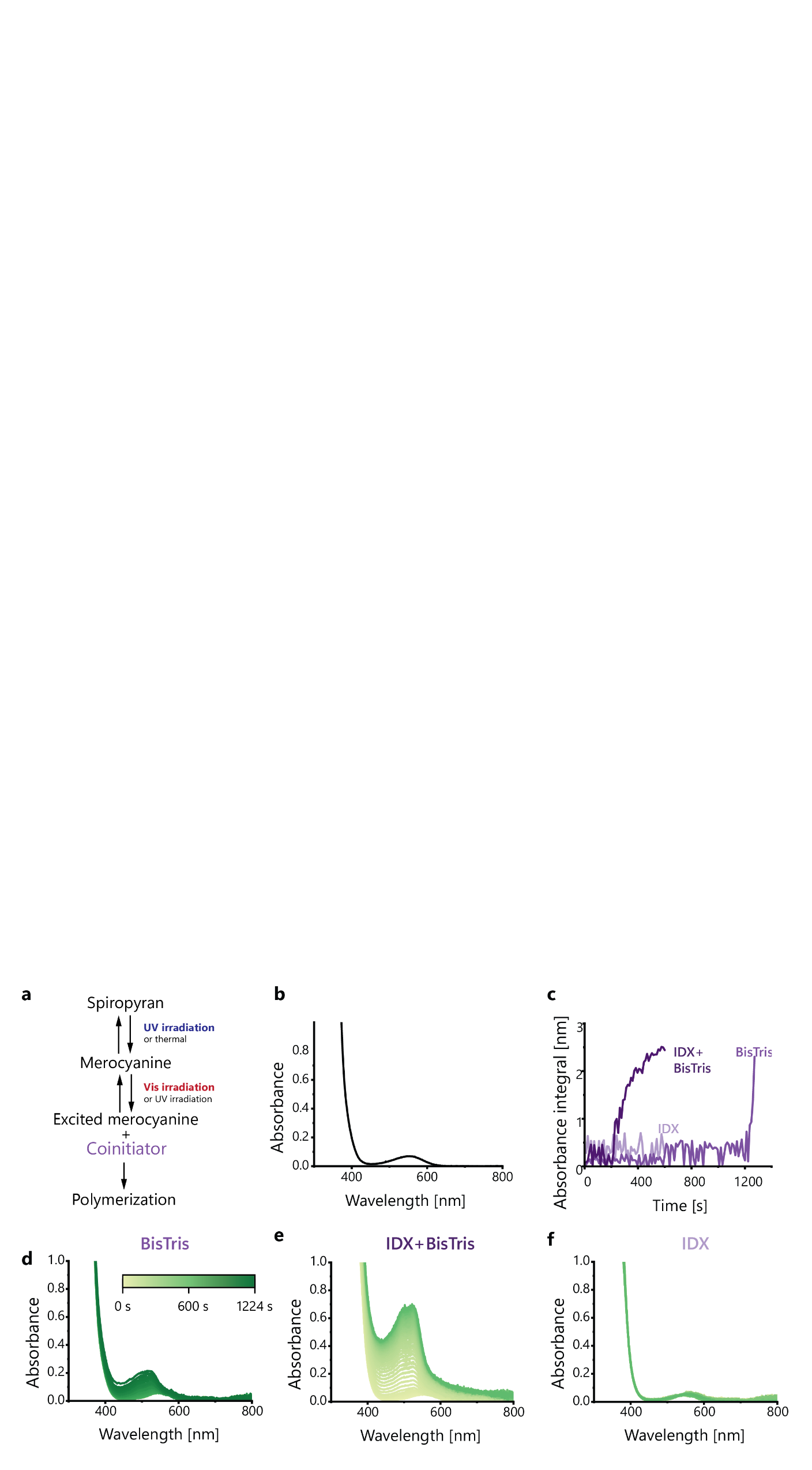

***Figure S3****. Effect of iodixanol on the dual-color photoinitiation process. (a) Schematic of the dual-color photoinitiation process. (b) Absorbance of the DCPI due to both the spiropyran (<450 nm) and merocyanine (450-650 nm) forms. (c) Absorbance integrated over time to show scattering due to PEGDA/acrylamide polymerization. (d) Absorbance from 0-1224 s, where LED intensity was increased from 80% to 100% at 600 s of formulations containing polymer, DCPI, and BisTris. (e) Absorbance from 0-600 s of formulations containing polymer, DCPI, BisTris, and IDX, which shows faster polymerization. (f) Absorbance from 0-600 s of formulations containing polymer, DCPI, and IDX, which show no polymerization.*

### Effect of printing components and light exposure on cell function

#### To determine the effect of printing components on cell function when exposed to light, 3T3 fibroblasts were exposed to iodixanol, DPI, and TEA at ideal print concentrations. When exposed to light, mitochondrial activity generally decreased and early apoptotic and dead cell fractions increased, regardless of which chemical the cells were exposed to (**Figure S4a-c**). Given the known detrimental effects of UV light on cell function, this is expected. At the optimized concentrations, there was no significant effect of chemical exposure on mitochondrial activity or early apoptotic cell fraction. However, cell death increased when exposed to both TEA and light. When exposed to increasing amounts of iodixanol, without light exposure mitochondrial activity increased and early apoptotic cell fraction decreased (**Figure S4d,e**). When irradiated with light, mitochondrial activity decreased and apoptosis increased at higher rates for higher concentrations of iodixanol. Importantly, at and below the concentration required for refractive index matching, there was no significant effect of iodixanol on mitochondrial activity or early apoptotic cell fraction. This confirms the results in **Figure 1d,** which suggests that iodixanol is not significantly cytotoxic, although it may have minor effects when irradiated with UV light. Regardless, when using iodixanol at its optimal dosage to normalize cell-induced light refraction that there are no detectable adverse effects, even when irradiated with UV light.

#### Methods

To determine effect of printing components under light exposure, monolayers of 3T3 fibroblasts were incubated with the desired component dissolved in PBS. Samples exposed to light were exposed to 10 mW/cm^2^ of UV light (Hamamatsu, Lightningcure) and 90 mW/cm^2^ visible light for five minutes, then incubated for an additional 25 minutes. Samples not exposed to light were placed directly in an incubator for 30 minutes. After incubation, samples were then washed twice with PBS, and new medium was added. 24 h later, the samples were stained with Hoechst 33342 (5 µg mL^-1^, Invitrogen), MitoTracker Green FM (100nM, Invitrogen), annexin V (Alexa Fluor 647 conjugate, 1:20, Invitrogen), and ethidium homodimer-1 (3 µm, Sigma-Aldrich) for 30 minutes. Samples were then imaged using a confocal laser scanning microscope (Zeiss 880 LSM) with a 20x Plan-APOCHROMAT objective. Images were segmented using the StarDist^[1]^ ImageJ plugin to find the total number of cells via Hoechst, the total number of dead or dying cells via ethD-1, and the total number of cells positive for annexin V. Cells considered early apoptotic were positive for annexin V and negative for ethD-1. In significance testing, means were compared within a given chemical concentration for light and dark and within light or dark between chemicals (e.g., TEA exposed to light was not compared to DPI kept dark).

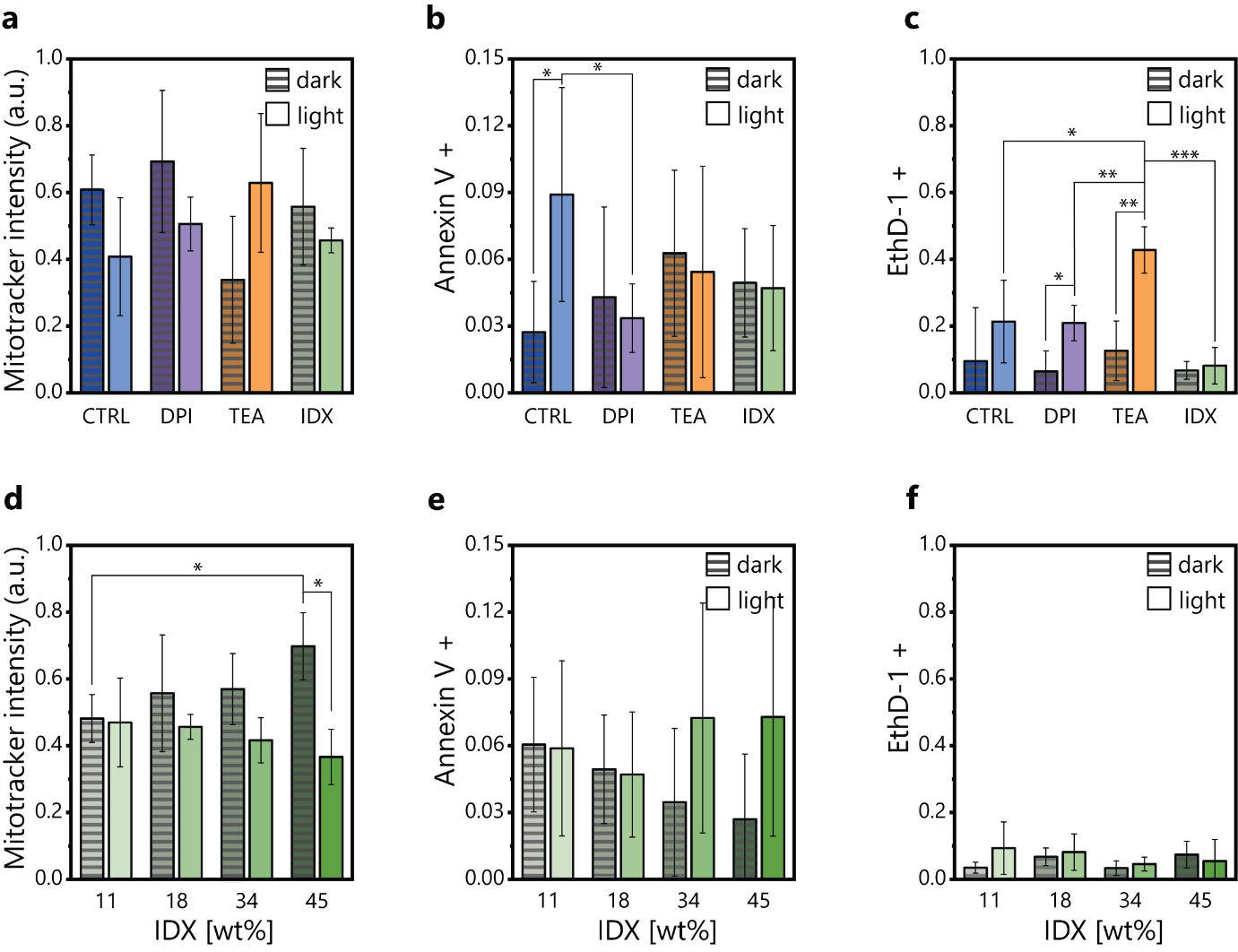

***Figure S4.*** *Effect of 0.03 wt% DPI, 1 wt% TEA, and 18% IDX on a monolayer of 3T3 fibroblasts exposed to UV and visible light or kept dark on (a) mitochondrial activity (b) early apoptotic cell fraction, and (c) dead cell fraction. Effect of iodixanol concentration on (d) mitochondrial activity, (e) early apoptotic cell fraction, and (f) dead cell fraction. All relationships not shown are not statistically significant. Analysis is a one-way ANOVA with a post-hoc Tukey test. *p<0.05, **p<0.01, ***p<0.001, n=3*

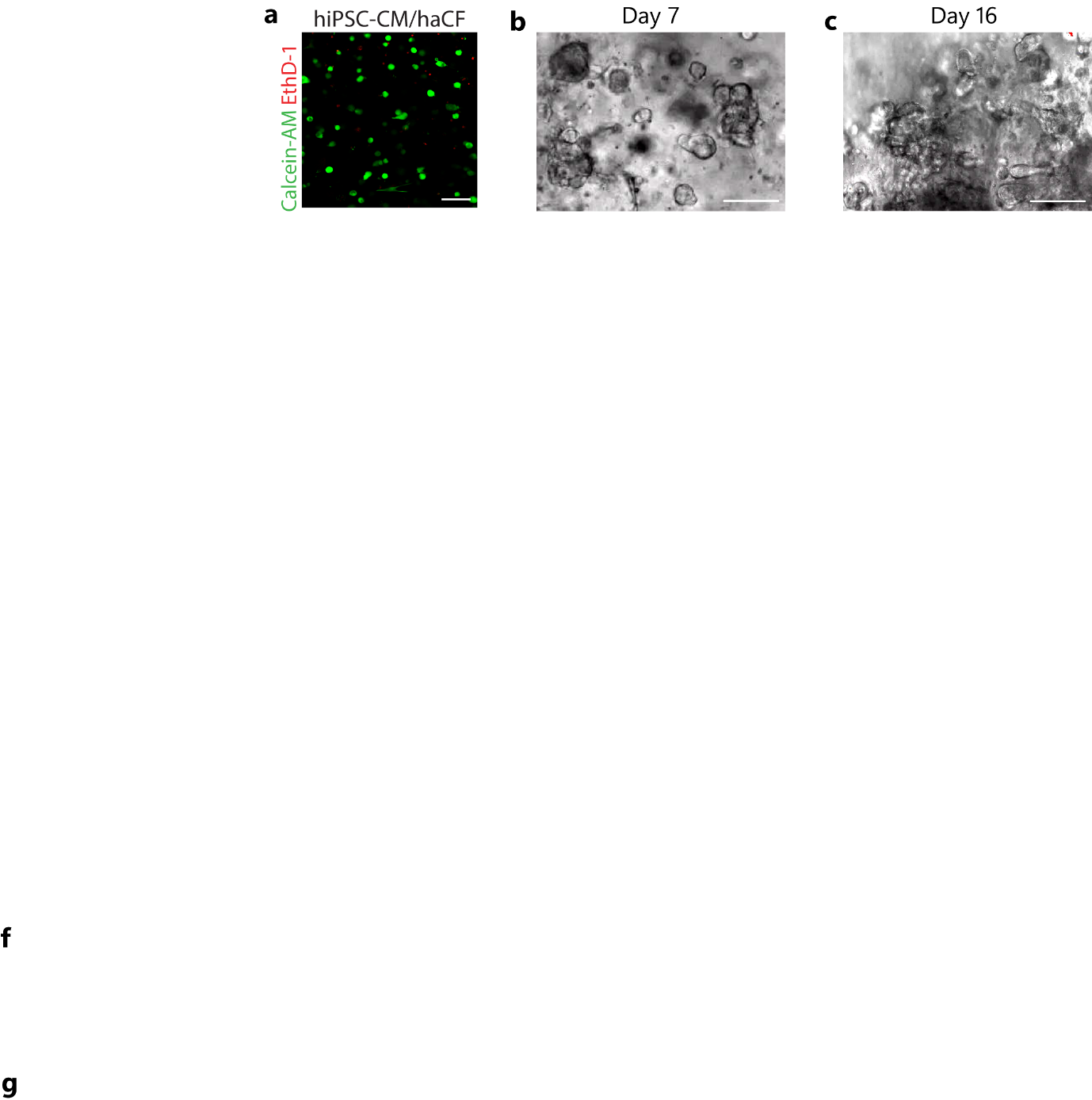

***Figure S5.*** *Xolographically printed constructs containing 10 M hiPSC-CMs mL^-1^ in 9% hPLMA and 1% GelMA. (a) Fluorescence micrographs of live-dead stained constructs containing hiPSC-CMs and haCFs 24 hours after Xolographic printing (scale bar: 50 µm). (b) Bright field images of printed constructs at days 7 and 16.*

**Movie S1.** Spontaneous contractions of hiPSC-CM-containing printed constructs in brightfield.

**Movie S2.** Spontaneous contraction of hiPSC-CM-containing printed constructs with calcium transient visualized.

**Movie S3.** Response to electrical pacing and pharmacological stimulation of constructs in 9% hPLMA and 1% GelMA with 10 M hiPSC-CM mL^-1^ initial cell density, in bright field and with calcium transient visualization.
